# Water and ion transport across the eversible vesicles in the collophore of a springtail *Orchesella cincta*

**DOI:** 10.1101/534164

**Authors:** Barbora Konopová, Dennis Kolosov, Michael J. O’Donnell

## Abstract

Springtails (Collembola) are ancient close relatives of the insects. The eversible vesicles are their unique paired transporting organs, which consist of an epithelium located inside a tube-like structure on the first abdominal segment called the collophore. The vesicles can be protruded out of the collophore and several lines of evidence indicate that they have a vital function in water uptake and ion balance. However, the amount of water absorbed by the vesicles and which other ions apart from sodium are transported remain unknown. Using *Orchesella cincta* as a model, we developed protocols for two assays that enabled us to study water and ion movement across the eversible vesicles in whole living springtails. Using an inverse Ramsay assay we demonstrate that the eversible vesicles absorb water from a droplet applied onto their surface. Using the scanning ion-selective electrode technique (SIET) we show that the vesicles absorb Na^+^ and Cl^−^ from the bathing medium, secrete NH_4_^+^, and both absorb and secrete K^+^, H^+^ is secreted at a low level in the anterior part and absorbed at the posterior. We did not detect transport of Ca^2+^ at significant levels. The highest flux was the absorption of Cl^−^, and the magnitude of ion fluxes were significantly lower in fully hydrated springtails. Our data demonstrate that the eversible vesicles are a transporting epithelium functioning in osmo- and ionoregulation, nitrogenous waste excretion and likely acid-base balance.

## INTRODUCTION

Springtails (Collembola) are a basal lineage of Hexapoda, which is a group of terrestrial arthropods that also includes the insects (Hopkin, 1997; Misof et al., 2014). The springtails are more primitive than the insects as they lack some of the advanced adaptations to terrestriality, including the Malpighian tubules for excretion and ion balance. Besides being important for understanding insect evolution, springtails also play a key role in the soil and stand as a model for ecotoxicology (Faddeeva-Vakhrusheva et al., 2016). Despite their importance, many basic aspects of springtail physiology remain unknown.

The hallmark of larval and adult springtails is the collophore (ventral tube) (Hoffmann, 1905; Imms, 1906), a tube-like structure in the centre of the ventral side of the first abdominal segment. The collophore originates from a pair of limbs that fuse together during embryogenesis (Imms, 1906; Uemiya and Ando, 1987; Konopova and Akam, 2014). The terminal part of the collophore is formed by the eversible vesicles, which are a special transporting epithelium. The vesicles are normally closed inside the tube, but can be everted by haemolymph pressure and retracted by muscles located inside the collophore (Eisenbeis, 1976).

Although the collophore is important for the springtails, its relevance to physiological processes of the animal remains unclear (Konopova and Akam, 2014; Favret et al., 2015). The organ is multifunctional, secreting, for instance, a substance with adhesive properties. However, most evidence to date indicates that its primary role is in uptake of water and transport of ions carried out by the eversible vesicles (e.g., Nutman, 1941; Noble-Nesbitt, 1963; Eisenbeis, 1982; Eisenbeis, 1974; Eisenbeis, 1976; Eisenbeis and Wichard, 1975a,b; Eisenbeis and Wichard, 1977).

The role in water balance has long been suspected (Noble-Nesbitt, 1963). When a dehydrated springtail encounters a puddle, the vesicles are everted and remain in contact with the water while the animal is drinking with its mouth. That fluid passes through the vesicles has been demonstrated by using vital dyes as indicators (Nutman, 1941). In the 1980s Eisenbeis and colleagues developed an assay that enabled estimation of the amount of water that is absorbed (Eisenbeis, 1982; Jaeger and Eisenbeis, 1984). Dehydrated springtails were allowed to walk on a wet filter paper and observed as they everted the vesicles. The rate at which water was taken up was calculated from the increase in animal weight after the experiment. These studies provided important evidence that the eversible vesicles are highly efficient in water uptake. Nevertheless, uptake by other means, such as the integument or mouth could not be easily controlled for in this assay. Thus, the evidence that significant amounts of water are taken up specifically by the eversible vesicles is still lacking.

Despite numerous clues that the eversible vesicles are ion-transporting organs, only the uptake of radioactively labeled sodium has been shown (Noble-Nesbitt 1963). A likely transport of chloride ion was indicated by histological staining with silver salts as markers (Eisenbeis and Wichard, 1975a,b). It is unknown if other ions are transported.

Here, we studied the springtail *Orchesella cincta*, which has previously been adapted as a model for ecotoxicology and developmental genetics (Konopova and Akam, 2014; Faddeeva-Vakhrusheva et al., 2018). We had two main goals. First, to establish a method for quantification of fluid transport specifically by the eversible vesicles. Second, to develop a protocol for electrophysiological measurement of ion transport across the eversible vesicles of *Orchesella* and to examine whether transport of selected ions by the vesicles indicates a role in osmo-and iono-regulation (Na^+^, K^+^, Cl^−^, Ca^2+^), nitrogenous waste excretion (NH_4_^+^) and acid-base balance (H^+^).

To quantitatively measure uptake of water by the eversible vesicles we adapted the classical Ramsay assay, which was developed for secretion of fluids by the insect Malpighian tubules (Ramsay, 1954). The original assay uses dissected tubules immersed in saline under paraffin oil and the rate of fluid secretion is calculated from the increasing size of a droplet secreted by the tubule. We show that an inverse Ramsay assay can be used for whole living *Orchesella*, in which changes in the size of a droplet in contact with the eversible vesicles are measured. Next, we present measurement of ion fluxes across the eversible vesicles using the scanning ion-selective electrode technique (SIET). We chose SIET because it enabled us to record ion transport rates in whole living springtails and on several different locations on the body in a single preparation. SIET uses ion-selective microelectrodes to measure ion concentrations in the measuring medium of the unstirred layer directly adjacent to the transporting cells and then further away (Piñeros et al., 1998; O’Donnell, 2009). From the differences in concentration of ions, fluxes are calculated, thus providing values for ion absorption (influx) or secretion (efflux). We show that the eversible vesicles absorb water from a droplet and transport a range of physiologically relevant ions. Our results demonstrate that the eversible vesicles are a multifunctional transporting epithelium and an important component of springtail osmoregulatory and excretory physiology.

## MATERIALS AND METHODS

### Animals

The culture of *Orchesella cincta* was maintained in Petri dishes with a base of plaster of Paris and fed with algae (*Pleurococcus* sp.) and yeast as described previously (Konopova and Akam, 2014). The animals were kept and experiments carried out at a laboratory temperature. Adults, mixed males and females, were used for all measurements. Animals designed for experiments were mildly dehydrated: they were transferred into a dry dish with food and kept there for 1-3 days.

### Inverse Ramsay assay

Adult springtails were briefly anesthetized with CO_2_ and using forceps immersed into heavy paraffin oil (Caledon Laboratory Chemicals, Georgetown, Canada) in a Sylgard coated dish. They were restrained between minutien pins to keep them in place and to isolate the collophore from moving appendages. Care was taken so that the integrity of the integument remained intact without haemolymph leaks. The collophore was brought into contact with a droplet of fluid whose composition mimicked ground water (modeled after Schouten and van der Brugge, 1989) (in mmol l^−1^): 0.1 KCl, 0.2 NaCl, 0.3 NH_4_Cl, 0.17 CaCl_2_, 0.15 MgSO_4_, 1.7 KNO_3_, 1 HEPES, pH=6.0). The diameter (d) of the 1 μl droplet that was pipetted onto the everted vesicles (experiment) or freely into the oil (control) was measured at 800x magnification using a calibrated ocular micrometer. The absorption rate (AR, nl min^−1^) by the vesicles was calculated from the change in droplet volume (4/3πr^3^, where r is the droplet radius) over 30 minutes. To be expressed per absorptive area, the value was divided by the absorptive area of the vesicles (0.110 mm^−2^), which was determined for *Orchesella* previously (Eisenbeis, 1982).

### SIET

The hardware, software and methodology for acquiring SIET data and calculating ion fluxes have been described previously (Donini and O’Donnell, 2005; O’Donnell and Ruiz-Sanchez, 2015; Kolosov et al., 2018a). Briefly, SIET measurements were made with hardware from Applicable Electronics (Forestdale, MA, USA) and Automated Scanning Electrode Technique (ASET) software (version 2.0; Science Wares, Falmouth, MA, USA). Micropipettes were pulled on a P-97 Flaming-Brown pipette puller (Sutter Instruments Co., Novato, CA, USA) from 1.5 mm borosilicate glass (World Precision Instruments Inc., Sarasota, FL, USA). At each measurement site, the ion-selective microelectrode was vibrated perpendicular to the tissue surface between two positions separated by 50 μm. The measured voltage gradients between the two points were converted into concentration gradients using the following equation

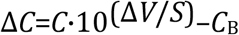

where Δ*C* is the concentration gradient between the two points measured in μmol cm^−3^; *C*_B_ is the background ion concentration, calculated as the average of the concentrations at each point measured in μmol l^−1^; Δ*V* is the voltage gradient obtained from ASET in μV; and *S* is the slope of the electrode.

Fluxes were estimated from the measured concentration gradients using Fick’s law:

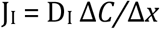

where J_I_ is the net flux of the ion in pmol cm^−2^ s^−1^; D_I_ is the diffusion coefficient of the ion (1.55×10^−5^ cm^2^ s^−1^ for Na^+^ and Cl^−^; 1.92×10^−5^ cm^2^ s^−1^ for K^+^; 1.19×10^−5^ cm^2^ s^−1^ for Ca^2+^, 9.4×10^−5^ cm^2^ s^−1^ for H^+^ and 2.09×10^−5^ cm^2^ s^−1^ for NH_4_^+^); ΔC is the concentration gradient in μmol cm^−3^; and Δx is the distance between the two points measured in cm. Microelectrodes were constructed with the following ionophores (Sigma-Aldrich, St. Louis, USA) (backfill and calibrating solutions (in mmol l^−1^) indicated in brackets): Na^+^ ionophore X (150 NaCl backfill, 0.15/1.5 NaCl calibrations); K^+^ ionophore I, cocktail B (150 KCl backfill, 1.5/15 KCl calibrations); Cl^−^ ionophore I, cocktail A (150 KCl backfill, 0.15/1.5 NaCl calibrations); Ca^2+^ ionophore I, cocktail A (100 CaCl_2_ backfill, 0.15/1.5 CaCl_2_ calibrations); H^+^ ionophore I, cocktail B (100 NaCl/100 Na-Citrate pH=6 backfill, 1 HEPES, pH=6/7 calibrations); NH_4_^+^ ionophore I, cocktail A (100 NH_4_Cl backfill, 0.15/1.5 NH_4_Cl calibrations). For measurements of Na^+^, K^+^ and Ca^2+^, the bathing medium was that described above for the inverse Ramsay assay. To avoid interference of competing ions with ionophores employed in SIET, the original recipe had to be modified for the measurement of NH_4_^+^ ions (0.1 mM KNO_3_ and 1.7 mM KCl were replaced with N-methyl-D-glucamine chloride to avoid K^+^ interference with NH_4_^+^ ionophore) and Cl^−^ ions (1.7 mM KNO_3_ was replaced with 0.85 mM K_2_SO_4_ to avoid NO_3_^−^ interference with Cl^−^ ionophore). H^+^ was measured in the medium adapted for NH_4_^+^ measurements.

The scanning was carried out as follows: each scan started at the anterior most part of one of the paired vesicles and continued at 25-μm intervals until the posterior most part. Protocol 1: If no regional signal heterogeneity was observed (Na^+^, Cl^−^, NH_4_^+^), three to five values surrounding the signal maximum were averaged and used as a representative n=1 for the biological replicate. Protocol 2: If anterior-posterior heterogeneity was observed (H^+^), three to five values surrounding signal maxima were averaged independently in the anterior and posterior lobes of the vesicle. Protocol 3: if anterior-posterior heterogeneity was present only in some of the samples, approaches 1 or 2 were followed depending on whether that particular sample exhibited the heterogeneity.

### Measurement of haemolymph ion concentration

Springtails were immersed in paraffin oil and one or two antennae were cut off at the midway point. Haemolymph was collected at the cut end of the antenna using a glass microcapillary pulled to a fine point. Samples were expelled under oil in a separate dish and analyzed with ion-selective microelectrodes as described previously (Naikkwah and O’Donnell, 2011; Kolosov et al., 2018b). Collecting samples from the antennae is a convenient method that provides relatively clean haemolymph as compared to sampling from around the collophore, which contains fat body. The haemolymph composition is expected to be similar in different parts of the body.

Preliminary results indicated that an unknown factor interfered with Na^+^ ionophore X-based electrodes in haemolymph samples as reported previously (Kolosov et al, 2018b). Therefore, Na^+^ ionophore III-based electrodes were used to avoid this. A solid-state Cl^−^ microelectrode was employed (construction described in Donini and O’Donnell, 2005) as Cl^−^ ionophore I ionophore is affected by other anions abundant in the haemolymph (e.g., HCO_3_^−^). The K^+^ and Ca^2+^ microelectrodes were based on the ionophores and backfill solutions described above for SIET. Calibration solutions were as follows: Na^+^ - 15mM and 150 mM NaCl; K^+^ - 1.5 mM and 15 mM KCl; Cl^−^ - 150 mM and 15 M KCl; Ca^2+^ 0.15 and 1.5 mM CaCl_2_; H^+^ - 1 mM HEPES pH=8 and pH=9.

### Statistics

Significant differences in (i) transport of Cl^−^ between fully hydrated and mildly dehydrated springtails and (ii) H^+^ flux between anterior and posterior collophore regions were determined using a Student’s t-test in SigmaPlot (version 11), with a P<0.05 limit.

### Image processing

Brightness and contrast in micrographs was adjusted using Adobe Photoshop CC 2017.1.1.

## RESULTS

### An inverse Ramsay assay quantifies the water absorbed specifically by the eversible vesicles of living *Orchesella*

The original Ramsay assay that we adapted here uses dissected organs immersed in paraffin oil. Our setup uses whole living adult *Orchesella* (Fig. 1A). Because the cuticle on the body of springtails is highly hydrophobic (Gundersen et al., 2017), the animals immerse easily into the oil. They survive the immersion well and stay alive for several hours under oil. After return to the culture they recover fully. Unlike the rest of the body, the cuticle on the eversible vesicles is hydrophilic and represents the only part of the animal that is wettable (Fig. 1B). The quantification of fluid absorption by the vesicles is based on measuring the decrease in size of a droplet that is dispensed onto their surface.

**Figure 1:**
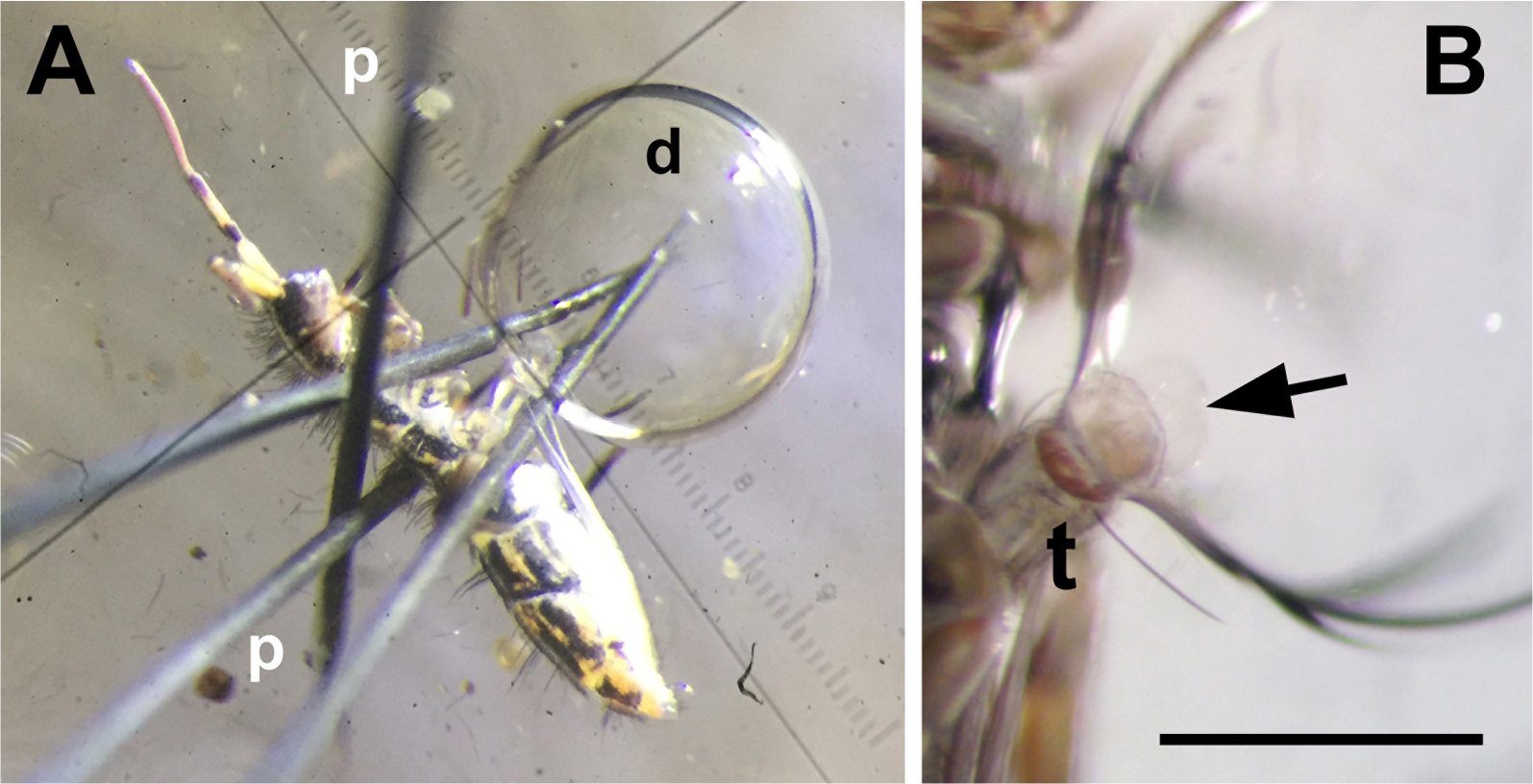
Measurement of water uptake by the eversible vesicles of *Orchesella* using an inverse Ramsay assay. Adult *Orchesella* was immersed in paraffin oil and clipped between minutien pins. A droplet of fluid was dispensed onto the everted vesicles. The springtails usually everted the vesicles under the oil themselves or were forced to do so by gentle pressure on the abdomen. The size of the droplet was measured using a calibrated ocular micrometer. The cuticle on the body of *Orchesella* is hydrophobic, thus the animals immersed easily into the oil and sank to the bottom of the dish. (B) The cuticle on the vesicles is hydrophilic and attached well to the droplet. The everted vesicles are marked with an arrow. Anterior is to the top left in (A) and to the top in (B). d, droplet of the measuring fluid; p, pins; t, tube of the collophore. Scale bar in (B), 0.5 mm.

To find out at what rate the eversible vesicles likely absorb water in nature we let them absorb a medium that mimics the composition of ground water (Schouten and van der Brugge, 1989) and which *Orchesella* is likely to encounter in its natural habitat. A 1 μl droplet of the medium was dispensed onto the everted vesicles of springtails that had been mildly dehydrated (see Materials and methods). Previous studies indicated that the main absorption period after the vesicles come into contact with water is 10-30 minutes (Eisenbeis, 1982). We monitored the change in size of the droplet after 30 minutes and from the values calculated a rate of fluid uptake 2.55±0.40 nl min^−1^ (N=8, s.e.m.). All springtails used in this experiment remained alive after they were removed from the oil. As a control, 1 μl droplets of the medium were kept under oil without contact with the springtail; none of the five droplets that we observed changed their size after 30 minutes. This demonstrates that the decrease in size detected in the experiment did not take place by diffusional loss of water into the oil. Taken together, the inverse Ramsay assay is a suitable method for precise measurement of the rate of fluid absorption by the eversible vesicles.

### SIET detects significant fluxes of Na^+^, Cl^−^, K^+^, H^+^ and NH_4_^+^ but not Ca^2+^ ions across the eversible vesicles of *Orchesella*

Animals were restrained for SIET as for the inverse Ramsay assay, but were bathed in medium mimicking ground water instead of paraffin oil. Because of their hydrophobic cuticle the springtails tend to float on the surface of the medium and were restrained with bent minutien pins as shown in Fig. 2A,A’). The springtails remained alive during the recording as they moved occasionally and fully recovered after the experiment.

**Figure 2:**
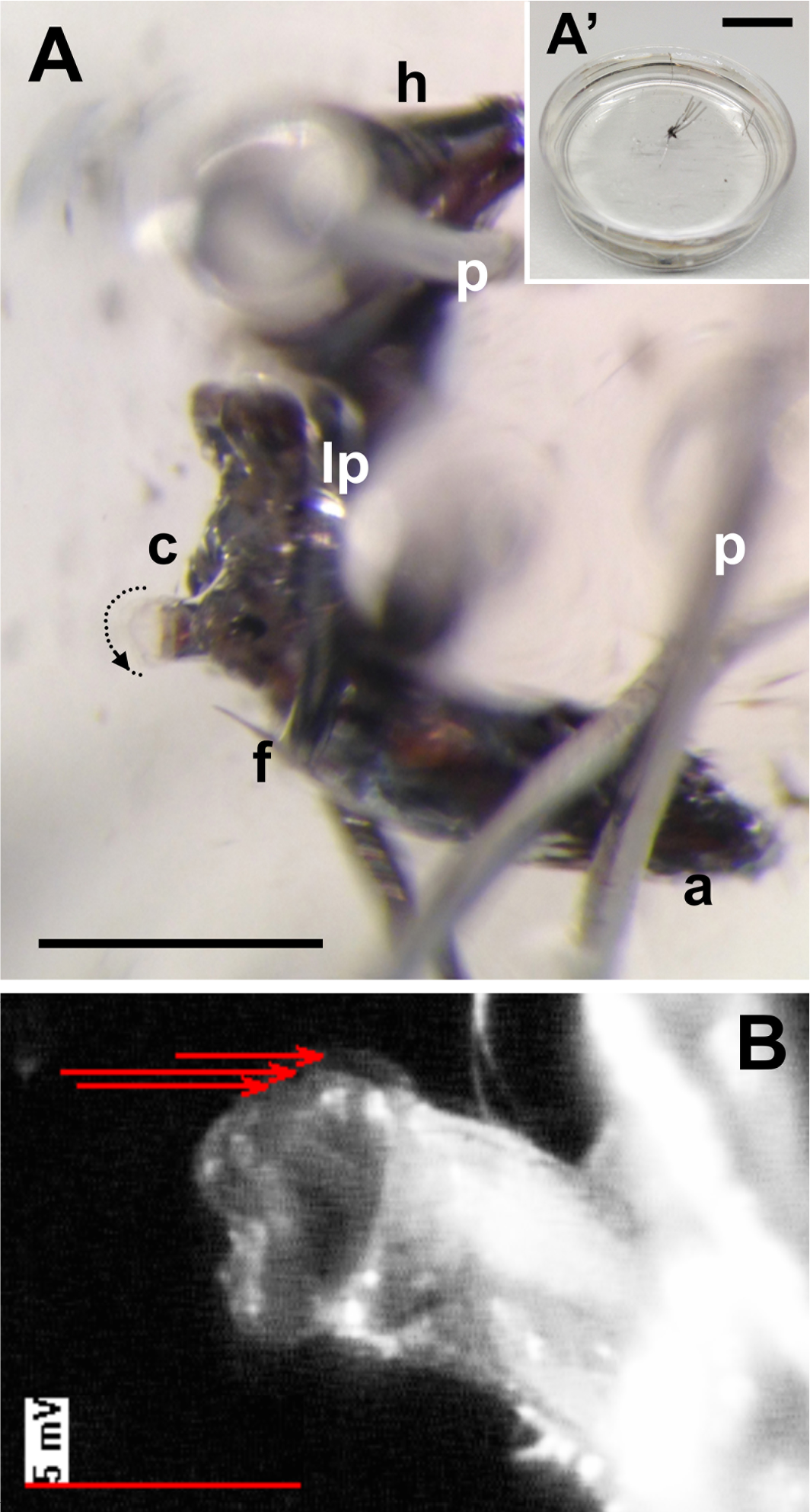
Preparation for SIET measurement of ion fluxes across the eversible vesicles of *Orchesella*. (A) Adult *Orchesella* were immersed into the measuring medium and carefully secured with minutien pins without the integument being pierced. The collophore protrudes through the loop made from a pin. Some individuals everted the vesicles themselves during mounting, others had to be gently pushed from the back. The animal in this photograph was immediately moving after it was returned to the culture; after a few minutes of rest it crawled away. The dotted line with an arrow around the vesicles indicates how the scanning was done from anterior to posterior. The inset (A’) shows the dish with the preparation. (B) Representative scan from SIET. Uptake of Cl^−^ in the anterior part of the eversible vesicles is shown. The length and direction of the arrows indicate the magnitude and direction of the flux, respectively. The magnitude is expressed as the difference in voltage between recording by the microelectrode close to the cells of the vesicles and then farther away (marker 5mV). The values were then used for calculation of concentration difference, which was converted into the net flux (pmol cm^−2^ s^−1^). Head to the top and ventral side to the left in all photographs. a, abdomen; c, collophore; f, furca; h, head; lp, pin making a loop; p, pins. Scale bars: in (A), 1 mm; in (A’), 1 cm.

We next used SIET to examine, if we could detect the flow of Na^+^, Cl^−^, K^+^, Ca^2+^, NH_4_^+^ and H^+^ across the vesicles (Fig. 2B). All our experimental animals were mildly dehydrated, as in the inverse Ramsay assay. The microelectrode recordings were carried out at several locations around the sphere of one of the everted vesicles from anterior to posterior (Fig. 2A). Vesicles remained everted during the whole time of recording, as monitored by a camera on the stereomicroscope that was attached to the SIET rig. Control recordings on two separate preparations of the collophore with retracted vesicles and on a special abdominal appendage called the furca that is used for jumping (Hopkin, 1997) did not detect any measurable Na^+^ flux. In contrast, significant fluxes of all examined ions except Ca^2+^ were detected across the vesicles (Fig. 3).

**Figure 3:**
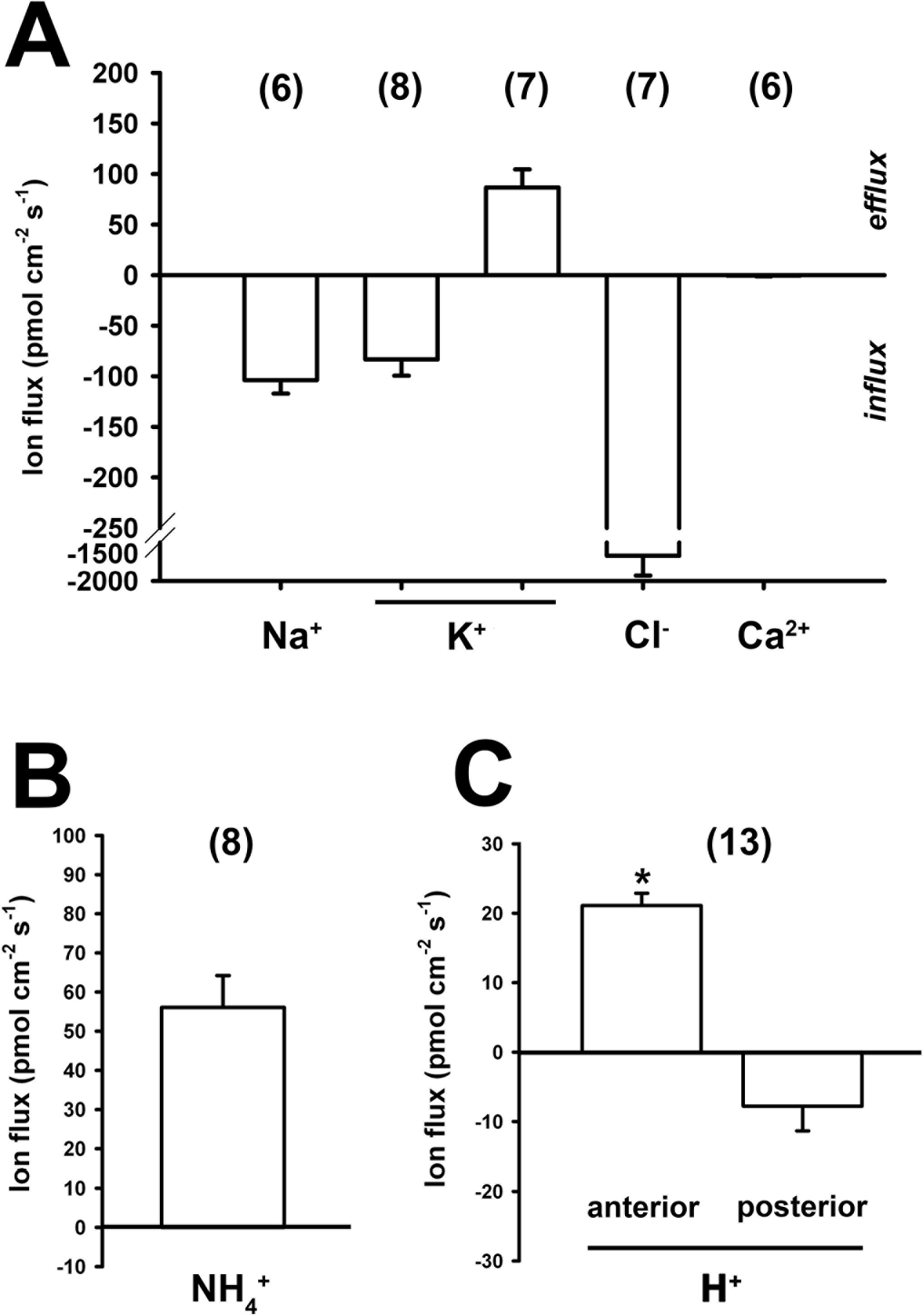
Ion fluxes across the eversible vesicles of *Orchesella* measured by SIET. Fluxes of ions implicated in osmo- and iono-regulation (A), nitrogenous waste excretion (B) and acid-base balance (C) were measured on mildly dehydrated springtails (see Materials and methods). The asterisk in (C) indicates significant difference between anterior and posterior as determined by a Student’s t-test. Values are mean ± s.e.m. Numbers in brackets above the bars indicate the number of animals examined (N). Negative values indicate ion uptake (influx), positive values indicate ion loss (efflux).

Na^+^ and Cl^−^ ions were absorbed by the vesicles of all animals and across all anterior to posterior locations on the vesicles that we examined. Mean values were −104 and −1541 pmol cm^−2^ s^−1^, for Na^+^ and Cl^−^, respectively (Fig. 3A). K^+^ ions were taken up by half of the animals, but secreted by others (Fig. 3A). Both influx and efflux of K^+^ occurred in all anterior to posterior locations on the vesicles, suggesting that unidirectional movement is not regionalized. The magnitude of 83±16 pmol cm^−2^ s^−1^, and efflux, 86±18 pmol cm^−2^ s^−1^, was similar. These results were obtained on animals examined simultaneously and kept under the same condition. The distribution of animals demonstrating the influx or efflux was random within the group. This might reflect as yet unknown endogenous physiological differences in K^+^ homeostasis between the animals. NH_4_^+^ ions were secreted by all animals at all anterior to posterior locations on the vesicles (Fig. 3B). The values were 56±8.1 pmol cm^−2^ s^−1^. H^+^ ions were secreted in the anterior half of the vesicles (efflux 21±1.8 pmol cm^−2^ s^−1^) and simultaneously absorbed (influx −7.8±3.6 pmol cm^−2^ s^−1^) across the posterior half (Fig. 3C) in all animals. In summary, these experiments demonstrate that the eversible vesicles transport a range of ions and the notably highest fluxes in our experiments were of Cl^−^.

### Uptake of Cl^−^ ions is lower in fully hydrated springtails

Next we examined whether the mild dehydrations of our experimental animals had any effect on the ion fluxes. As an example we chose the Cl^−^ ion, for which we recorded the highest flux across the vesicles, and compared the flux between fully hydrated springtails and those that were used in the rest of our experiments. We found that the eversible vesicles of fully hydrated *Orchesella* still absorbed Cl^−^ ion, but this influx was ~13 times (significantly) lower (Fig. 4). Thus, mild dehydration of the whole animal leads to more pronounced fluxes of Cl^−^ across the eversible vesicles.

**Figure 4:**
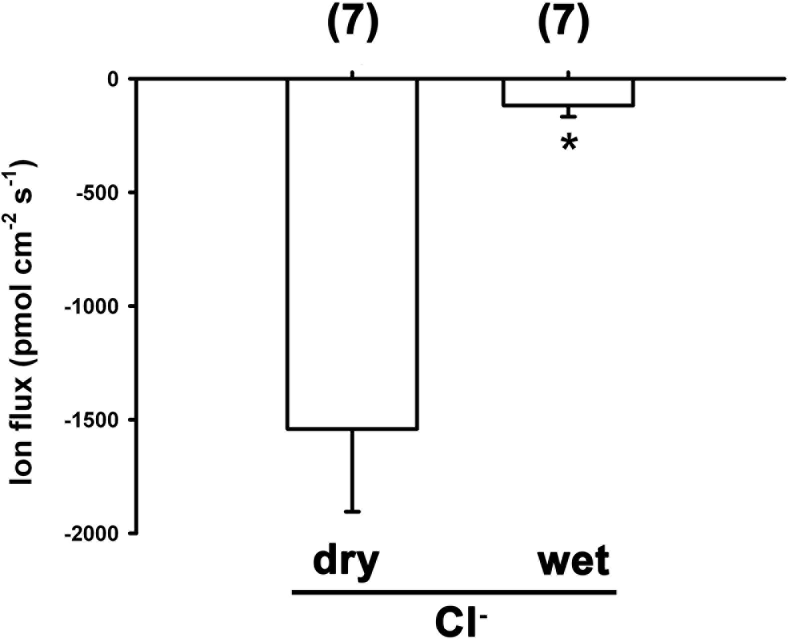
The level of *Orchesella* hydration affects the amount of Cl^−^ uptake by the eversible vesicles. The springtails kept at optimal humid conditions (“wet”) show significantly lower uptake of Cl^−^ by their eversible vesicles compared to animals previously exposed to mild dehydration (“dry”). The data for mildly dehydrated animals are from Fig. 3A. Values are mean ± s.e.m. Numbers in brackets above the bars indicate the number of animals examined (N). Asterisk indicates significant difference between dry and wet as determined by a Student’s t-test.

### Haemolymph concentrations of Na^+^, Cl^−^ and K^+^ indicate that the eversible vesicles transport these ions against the concentration gradient

Finally, we determined whether the exchange of ions between the eversible vesicles and external medium takes place against the concentration gradient, thus likely by an active transport. We compared the concentrations of Na^+^, Cl^−^ and K^+^ in the haemolymph and in the medium. Haemolymph concentrations were measured using ion-selective microelectrodes and concentrations in the medium were calculated from the recipe. The concentrations of Na^+^, Cl^−^ and K^+^ in *Orchesella* haemolymph were measured previously (Klein et al., 2008) and the values that we obtained for our springtails were 0.86, 1.56 and 0.85 times different, respectively. The concentration of all these three ions in the haemolymph is higher than in the measuring media (Table 1). This indicates that the uptake of such ions takes place against a concentration gradient.

**Table 1.** Comparison of ion concentration in the haemolymph and the measuring medium. Haemolymph ion concentrations were measured with ion-selective microelectrodes. Values are mean ± s.e.m. (N). Ion concentrations in the measuring medium were calculated from the recipe.

| Ion | Concentration (mM) |  |
| --- | --- | --- |
|  | Haemolymph | Measuring medium (theoretical) |
| Na<sup>+</sup> | 131.8 ± 5.3 (16) | 0.2 |
| Cl<sup>-</sup> | 178.3 ± 6.9 (13) | 0.94 |
| K<sup>+</sup> | 5.9 ± 0.7 (21) | 1.8 |

## DISCUSSION

### Eversible vesicles in the collophore of *Orchesella* are potent water absorbing organs

By employing the inverse Ramsay assay we found that the pair of eversible vesicles of mildly dehydrated *Orchesella* absorbs water at a rate 2.55 nl min^−1^. We recalculated this value per absorptive area of the vesicles and obtained the rate 23.18 nl mm^−2^ min^−1^ (see Materials and Methods). How does this value compare (1) to the absorption rate measured on springtails previously and (2) to the transport rate across fluid transporting epithelia of insects?

Previous quantification of water absorption by eversible vesicles of *Orchesella* and closely related *Tomocerus* was based on weighing springtails before and after they were allowed to absorb fluid from a wet filter paper (Eisenbeis, 1982; Jaeger and Eisenbeis, 1984). The absorption rate, given by weight gain over a period of time, was expressed per mm^2^ of absorptive area of the vesicles. The rates from the filter paper absorption-weighing assay for *Orchesella* were 8 times higher for distilled water and 5 times higher for 25 mM NaCl (solutions with lower and higher osmolarity than the medium used in the present experiments) compared to our results. Our absorption rate is closest to the minima previously obtained for 25 mM NaCl (Eisenbeis, 1982). Why are the results from the two assays different? While the assay introduced by Eisenbeis and colleagues simulates natural conditions better, our inverse Ramsay assay is specific. It measures only the fluid taken up by the eversible vesicles and excludes drinking and absorption by other tissues. Other experimental conditions and the level of dehydration may differ.

How does the rate of absorption by the eversible vesicles compare to transport by other epithelia? The Ramsay assay on tissue immersed in paraffin was used for measurement of the rate of secretion from the Malpighian tubules. A single Malpighian tubule has a basal (unstimulated) secretory rate of typically less than 1 nl min^−1^ (e.g., Ramsay, 1954; Dow et al., 1994; Beyenbach, 2003), but up to 15 nl min^−1^ in some species (Kolosov et al., 2018a) (compare with absorption by the eversible vesicles 2.55 nl min^−1^). Expressed per size of secretion area the maximal rate of secretion by the Malpighian tubules measured on a wide range of insects is about 0.05-13 nl mm^−2^ min^−1^ (Philips, 1981), therefore typically less than half the absorption rate for the eversible vesicles of *Orchesella* measured in the current study (23.18 nl mm^−2^ min^−1^). The Ramsay assay was also used to quantify secretion by the “fastest fluid-secreting cell known”, the cells in the Malpighian tubules of *Rhodnius prolixus* (Maddrell, 1991). After the blood-meal, the secretion rate increases up to 46 nl mm^−2^ min^−1^ (expressed per the area of the secretory part) (Bradley, 1983). Based on these approximate comparisons it appears that the eversible vesicles of *Orchesella* transport water at a speed approximately half that that seen in the tubules of a blood-feeding hemipteran specifically adapted for rapid fluid secretion.

### The eversible vesicles transport ions at levels comparable to other transporting epithelia of insects

Our microelectrode recordings across the eversible vesicles using SIET detected significant fluxes of Na^+^, K^+^, Cl^−^, NH_4_^+^ and H^+^. In insect transporting epithelia the magnitudes of fluxes measured by ion-selective electrodes typically range 10-1000 pmol cm^−2^ s^−1^ (e.g., Donini and O’Donnell, 2005; Nguyen and Donini, 2010; Naikkhwah and O’Donnell, 2012; Pacey and O’Donnell, 2014; Paluzzi et al., 2014; Robertson et al., 2014; O’Donnell and Ruiz-Sanchez, 2015; D’Silva et al., 2017; Kolosov et al., 2018a). The fluxes that we obtained for Na^+^, Cl^−^, K^+^ and NH_4_^+^ (−104, −1541, −83/+86, +56 pmol cm^−2^ s^−1^, respectively; Fig. 3A,B) are of comparable magnitude. The influx of Na^+^ recorded is consistent with previous observations showing that the eversible vesicles pick up radiolabelled Na^+^ from ground water (Noble-Nesbitt, 1963). The influx of Cl^−^ is particularly remarkable. For example, the anal papillae of mosquitoes absorb Cl^−^ *in vivo* at a rate of 230 pmol cm^−2^ s^−1^ (Donini and O’Donnell, 2005).

### The eversible vesicles function in osmo- and ionoregulation, excretion and likely acid-base balance

Our SIET data provide support for the function of the eversible vesicles in osmo- and ionoregulation. Na^+^, Cl^−^ and K^+^ are among the major ions in the haemolymph. These were the most intensively transported ions in our mini screen (Fig. 3A). We suggest that when the springtail encounters a puddle of water, it uses the eversible vesicles to re-fill ions and maintain internal homeostasis (Fig. 5). The uptake of Na^+^, Cl^−^ and K^+^ likely takes place by an active transport, because their concentrations that we measured in the haemolymph are higher than in the external medium (Table 1). The active uptake of Na^+^, Cl^−^ and K^+^ then probably osmotically draws water into the cells (Maddrell, 1969). This may also bring in small organic molecules, such as urea or glycerol (Schreiber and Eisenbeis, 1985).

**Figure 5:**
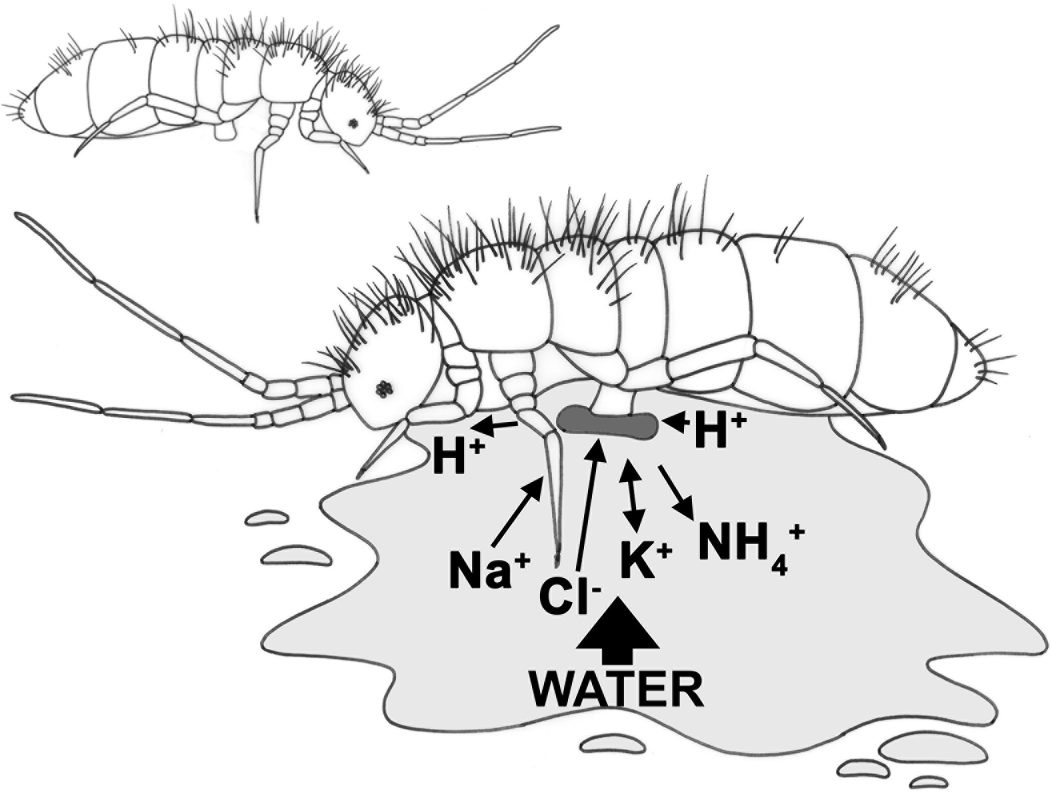
How the eversible vesicles of springtails are used in the nature. Springtails on dry surface (top) keep their eversible vesicles inside the tube of the collophore. When the springtail encounters water (bottom), the vesicles are everted. Water is absorbed and ions are exchanged: Na^+^, Cl^−^ are taken up, K^+^ taken up or secreted, NH_4_^+^ secreted, H^+^ secreted at the anterior part of the vesicles and absorbed at the posterior part. The length of the arrow corresponds approximately to the magnitude of the ion flux.

An observation not anticipated from previous studies was the secretion of NH_4_^+^ (Fig. 3B). This suggests that the eversible vesicles function also in nitrogen (metabolic) waste excretion. Generally, terrestrial insects excrete nitrogen waste as urea or uric acid, and aquatic insects excrete primarily NH_3_/NH_4_^+^, collectively known as ammonia (O’Donnell and Donini, 2017). While excreting ammonia is energetically advantageous, accumulation of this molecule in tissues is toxic. In aquatic animals, ammonia is carried away by the surrounding water. Perhaps the springtails use the opportunity to release NH_4_^+^ into the water puddle while absorbing water (Fig. 5).

Finally, we found that the eversible vesicles transport H^+^ ions, although at relatively low levels (Fig. 3C and discussed above). This result suggests that the epithelium might also function in regulation of acid-base homeostasis (pH), similar to the anal papillae of mosquito larvae (Donini and O’Donnell, 2005). We showed that H^+^ is secreted in the anterior and, at a lower magnitude of flux, taken up in the posterior. The difference in direction of the flux suggests that there is a functional specialization between the anterior and posterior part of the vesicles. It is worth noting that H^+^-selective microelectrode will detect transport of any ion that can buffer protons, such as HCO_3_^−^. Thus, the detected regional heterogeneity of H^+^ transport may indicate transport of different ions. It is also worth noting that the anterior half of the collophore and the vesicles contain the ventral groove (linea ventralis). This groove in the cuticle extends from the labial glands in the head and contains the products of the glands, called the urine (Hoffmann, 1905; Verhoef et al., 1979, 1983). Possibly, the urine in the anterior of the vesicles influences H^+^ concentrations.

### The eversible vesicles in the collophore are an important part of springtail physiology

The eversible vesicles are a multifunctional epithelium (Imms, 1906; Hopkin, 1997). The absorption of water is clearly important as it helps to speed up re-hydration of the animals living in the soil where water content might be changeable. Arctic and Antarctic springtails use dehydration as a strategy to survive extremely low temperature (Cannon and Block, 1988; Worland et al., 2010; Sørensen and Holmstrup, 2011). When favorable conditions return, and the springtails need to intake an amount of water, they “drink” with both their mouth and the vesicles (William Block, British Antarctic Survey, personal communication).

A peculiar part of springtail physiology is the absence of the Malpighian tubules (Humbert, 1974), a key insect organ for osmo- and ionoregulation, acid-base balance and excretion. How can springtails cope without them? In insects the Malpighian tubules (secretory epithelium) together with the hindgut (absorptive epithelium) make the “functional kidney”. While the springtails excrete nitrogen waste by accumulating it in the midgut epithelium that is removed at each moult (springtails continue moulting as adults) (Humbert, 1974, 1978), clues exist that the labial glands and the eversible vesicles form the “kidney” (Verhoef et al., 1979, 1983). Whether the secretion from the labial glands is re-absorbed at the eversible vesicles (Verhoef et al., 1983) or what actually occurs is unclear. This will be the subject of future research.

## ACKNOWLEDGEMENTS

Data were collected during BK’s visit to the lab of MJO; she acknowledges all members for hosting. BK thanks Simon Maddrell for a discussion on the project and his suggestion of the inverse Ramsay assay. We thank to Barry Denholm and William Block for comments on the draft.

## AUTHOR CONTRIBUTIONS

Conceptualization: BK; Methodology: BK, DK, MJO; Validation: BK, DK; Formal analysis: DK; Investigation: BK, DK, MJO; Resources: BK, DK, MJO; Writing – original draft: BK; Writing – review & editing: BK, DK, MJO; Visualization: BK, DK; Supervision: MJO; Project administration: BK, DK, MJO; Funding acquisition: MJO.

